# Characterization of the growth behavior of *Listeria monocytogenes* in *Listeria* synthetic media

**DOI:** 10.1101/2023.02.21.529469

**Authors:** Lisa Maria Schulz, Alicia Konrath, Jeanine Rismondo

## Abstract

The foodborne pathogen *Listeria monocytogenes* can grow in a wide range of environmental conditions. For the study of the physiology of this organism, several chemically defined media have been developed over the past decades. Here, we examined the ability of *L. monocytogenes* wildtype strains EGD-e and 10403S to grow under salt and pH stress in *Listeria* synthetic medium (LSM). Furthermore, we determined that a wide range of carbon sources could support growth of both wildtype strains in LSM. However, for hexose phosphate sugars such as glucose-1-phosphate, both *L. monocytogenes* strains need to be pre-grown under conditions, where the major virulence regulator PrfA is active. In addition, growth of both *L. monocytogenes* strains was observed when LSM was supplemented with the amino acid sugar *N*-acetylmannosamine (Man*N*Ac). We were able to show that some of the proteins encoded in the operon *lmo2795*-*nanE*, such as the Man*N*Ac-6-phosphate epimerase NanE, are required for growth in presence of Man*N*Ac. The first gene of the operon, *lmo2795,* encodes a transcriptional regulator of the RpiR family. Using electrophoretic mobility shift assays and quantitative real time PCR analysis, we were able to show that Lmo2795 binds to the promoter region of the operon *lmo2795*-*nanE* and activates its expression.

**Originality-Significance Statement:** Current knowledge of growth and survival of the human pathogen *Listeria monocytogenes* under diverse stress conditions is mostly generated in complex medium, a condition that is rarely found in the environment or host of the pathogen. Our work contributes to the characterization of the physiology of *L. monocytogenes* grown under nutrient limiting conditions and its growth requirements with regards to metabolizable carbon sources.

## Introduction

The foodborne pathogen *Listeria monocytogenes* is a Gram-positive, facultative anaerobic, rod-shaped bacterium, which is ubiquitously found in nature. It can survive as a saprophyte on decaying plant material, in soil or in wastewater. *L. monocytogenes* and other *Listeria* species can also enter food processing facilities and food chains, for instance due to contaminated water or raw materials (Lourenco *et al*., 2022). Several recent outbreaks of *L. monocytogenes* were associated with contaminated fruits and vegetables, such as melons, apples or mushrooms, or contaminated meat products (Matle *et al*., 2020; Lachmann *et al*., 2021; Townsend *et al*., 2021). The intracellular pathogen *L. monocytogenes* is the causative agent of the disease listeriosis. In healthy individuals, the intake of *L. monocytogenes* via the consumption of contaminated food products mostly leads to mild symptoms, such as fever and a self-limiting gastroenteritis. However, for immunocompromised individuals, the elderly, newborns and pregnant women, the infection with *L. monocytogenes* can cause more severe symptoms, such as muscle aches, vomiting, diarrhea, encephalitis or meningitis. For pregnant women, it can also lead to premature or still birth (Radoshevich and Cossart, 2018). This is achieved by its ability to cross all human barriers, namely the intestinal, placental and blood-brain barriers (Lecuit, 2005).

The survival of *L. monocytogenes* in food processing facilities and within the host is enabled due to its ability to grow under diverse stress conditions (reviewed in (Gaballa *et al*., 2019; Wiktorczyk-Kapischke *et al*., 2021; Osek *et al*., 2022)). Previous studies reported that *L. monocytogenes* can grow at temperatures between 1 and 45°C, in a pH range from 4.0 to 9.5 or in the presence of up to 10% salt (Patchett *et al*., 1992; Arizcun *et al*., 1998; Vasseur *et al*., 2001). *L. monocytogenes* is also able to use a variety of carbon sources ranging from simple monosaccharides (glucose, fructose) to more complex sugars (maltodextrin) and polyols (glycerol, D-arabitol, D-xylitol) (Gopal *et al*., 2010; Deutscher *et al*., 2014; Kentache *et al*., 2016). The ability to take up such a wide range of carbon sources is facilitated by a diverse set of transporters, which are encoded in the genome of *L. monocytogenes*. Most of the carbon sources are imported into the cell via the phosphoenolpyruvate:carbohydrate phosphotransferase system (PTS). The genome of *L. monocytogenes* wildtype strain EGD-e contains 86 PTS genes, which encode for 29 complete and 12 incomplete PTS complexes (Glaser *et al*., 2001; Stoll and Goebel, 2010). These PTS transporters are involved in the transport of diverse carbon sources such as glucose, fructose, D-arabitol and D-xylitol (Deutscher *et al*., 2014; Kentache *et al*., 2016). Some of these PTS transporters have also overlapping substrate specificities. For instance, the two PTS systems ManLMN (MptACD) and MpoABCD, encoded by *lmo0096*-*0098* and *lmo0784*-*0781*, respectively, are both involved in the transport of glucose and mannose (Stoll and Goebel, 2010; Aké *et al*., 2011). Inactivation of all PTS systems, e.g. by deleting the phosphotransferase system enzyme I encoding gene *ptsI*, did not abolish fermentation of some sugars such as glucose, fructose or mannose, suggesting that *L. monocytogenes* possesses also other sugar transporters (Deutscher *et al*., 2014). Indeed, *L. monocytogenes* encodes three GlcU-like non-PTS permeases (Lmo0169, Lmo0176, Lmo0424), which could be involved in the import of these sugars (Aké *et al*., 2011; Deutscher *et al*., 2014). The import of L-rhamnose is accomplished by the major facilitator superfamily protein Lmo2850, which uses a proton symport mechanism (Fieseler *et al*., 2012; Deutscher *et al*., 2014; Zeng *et al*., 2021). Within the host, *L. monocytogenes* uses mainly glucose-1-phosphate, glucose-6-phosphate and glycerol as carbon sources (Ripio *et al*., 1997; Grubmüller *et al*., 2014). The transporter Hpt, also named UhpT, takes up hexose phosphates from the host cell cytosol (Chico-Calero *et al*., 2002), while the glycerol uptake facilitator GlpF facilitates the import of glycerol (Joseph *et al*., 2008). The expression of *hpt* thereby depends on PrfA, the main virulence regulator of *L. monocytogenes* (Ripio *et al*., 1997; Chico-Calero *et al*., 2002). *L. monocytogenes* possesses eight predicted carbohydrate ATP-binding cassette (ABC) transporters. One of these transporters, which is encoded by *lmo2123-2125*, is required for the import of maltose and maltodextrin (Gopal *et al*., 2010). The remaining seven ABC transporters have not yet been studied.

For many studies on the physiology and stress tolerance of *L. monocytogenes*, bacteria were cultured in complex medium such as brain heart infusion (BHI) or tryptic soy broth (TSB). In their natural habitat or within their host, *L. monocytogenes* likely encounters more limiting growth conditions. Within the last decades, several minimal media have been developed which can be used to grow *L. monocytogenes* (Welshimer, 1963; Premaratne *et al*., 1991; Phan-Thanh and Gormon, 1997; Tsai and Hodgson, 2003; Jarvis *et al*., 2016; Whiteley *et al*., 2017). In this study, we aimed to characterize the growth of the two widely used *L. monocytogenes* wildtype strains EGD-e and 10403S, which belong to serotype 1/2a, in the recently developed *Listeria* synthetic medium (LSM) (Whiteley *et al*., 2017). *L. monocytogenes* strains belonging to serotype 1/2a are often recovered from food, environmental and clinical samples (Ward *et al*., 2008; Orsi *et al*., 2011) and thus encounter diverse environmental conditions. We thus determined the growth behavior of EGD-e and 10403S in LSM under stress conditions, which they encounter in food or the environment or within the host during infection, namely high salt and low pH stress. To our knowledge, growth of *L. monocytogenes* in LSM was so far only assessed with glucose as a sole carbon source. We therefore determined, which other carbon sources can facilitate growth of *L. monocytogenes* EGD-e and 10403S in LSM. While 95% of the open reading frames are conserved between EGD-e and 10403S, the genomes differ by over 30,000 single nucleotide polymorphisms (Glaser *et al*., 2001; Bécavin *et al*., 2014). It is thus not surprising that we could observe differences in the growth behavior between these two strains.

## Experimental procedures

### Bacterial strains and growth conditions

All strains and plasmids used in this study are listed in Table 1. *Escherichia coli* strains were grown in Luria-Bertani (LB) medium and *Listeria monocytogenes* strains in brain heart infusion (BHI) medium or *Listeria* synthetic medium (LSM) at 37°C unless otherwise stated. Where required, antibiotics and supplements were added to the medium at the following concentrations: for *E. coli* cultures, ampicillin (Amp) at 100 µg ml^-1^, kanamycin (Kan) at 50 µg ml^-1^, and for *L. monocytogenes* cultures, chloramphenicol (Cam) at 7.5 µg ml^-1^ and Kan at 50 µg ml^-1^.

**Table 1:** Bacterial strains used in this study

| Unique ID | Strain name and resistance | Source |
| --- | --- | --- |
| ***Escherichia coli* strains** |  |  |
| ANG1264 | DH5α pKSV7; AmpR | (Smith and Youngman, 1992) |
| EJR149 | XL10 Gold pWH844; AmpR | (Schirmer *et al.*, 1997) |
| EJR88 | XL1-Blue pKSV7-Δ*lmo*2795; AmpR | This study |
| EJR121 | XL1-Blue pKSV7-Δ*lmo*2800; AmpR | This study |
| EJR122 | XL10-Gold pKSV7-Δ*lmo*2799; AmpR | This study |
| EJR123 | XL10-Gold pKSV7-Δ*lmo*2798; AmpR | This study |
| EJR131 | DH5α pWH844-*lmo*2795; AmpR | This study |
| EJR164 | XL1-Blue pKSV7-Δ*lmo*2797; AmpR | This study |
| EJR210 | XL10-Gold pKSV7-Δ*nanE*; AmpR | This study |
| EJR236 | XL10-Gold pKSV7-Δ*lmo*2796; AmpR | This study |
| ***Listeria monocytogenes* strains** |  |  |
| ANG873 | EGD-e | (Glaser *et al.*, 2001) |
| ANG1263 | 10403S; StrepR | (Bishop and Hinrichs, 1987) |
| BUG2214 | EGD-e Δ*prfA* | (Mandin *et al.*, 2007) |
| LJR123 | EGD-e Δ*lmo*2795 | This study |
| LJR132 | EGD-e Δ*lmo*2798 | This study |
| LJR135 | EGD-e Δ*lmo*2799 | This study |
| LJR204 | EGD-e Δ*lmo*2797 | This study |
| LJR205 | EGD-e Δ*lmo*2799 Δ*lmo*2797 | This study |
| LJR206 | EGD-e Δ*lmo*2800 | This study |
| LJR207 | EGD-e Δ*nanE* | This study |

LSM was prepared in accordance with the recipe from Whiteley et al. with minor modifications (Whiteley *et al*., 2017). L-leucine, L-methionine and L-valine were used instead of the corresponding DL-enantiomers. L-arginine·HCl, riboflavin, Na_2_HPO_4_·7 H_2_O, CoCl_2_·7 H_2_O and L-cysteine·2·HCl were substituted for L-arginine, riboflavin 5’ monophosphate sodium salt hydrate, Na_2_HPO_4_·12 H_2_O, CoCl_2_·6 H_2_O and L-cysteine·HCl·H_2_O, respectively (see Table 2 for detailed LSM composition and preparation). The pH of LSM was adjusted with HCl, where indicated. Sodium chloride was added to LSM for characterization of the growth of *L. monocytogenes* strains under salt stress at the following concentrations: 1% (0.17 M), 2% (0.34 M), 3% (0.51 M), 4% (0.68 M), 5% (0.86 M) and 6% (1.03 M). To test the ability of *L*.

**Table 2:**
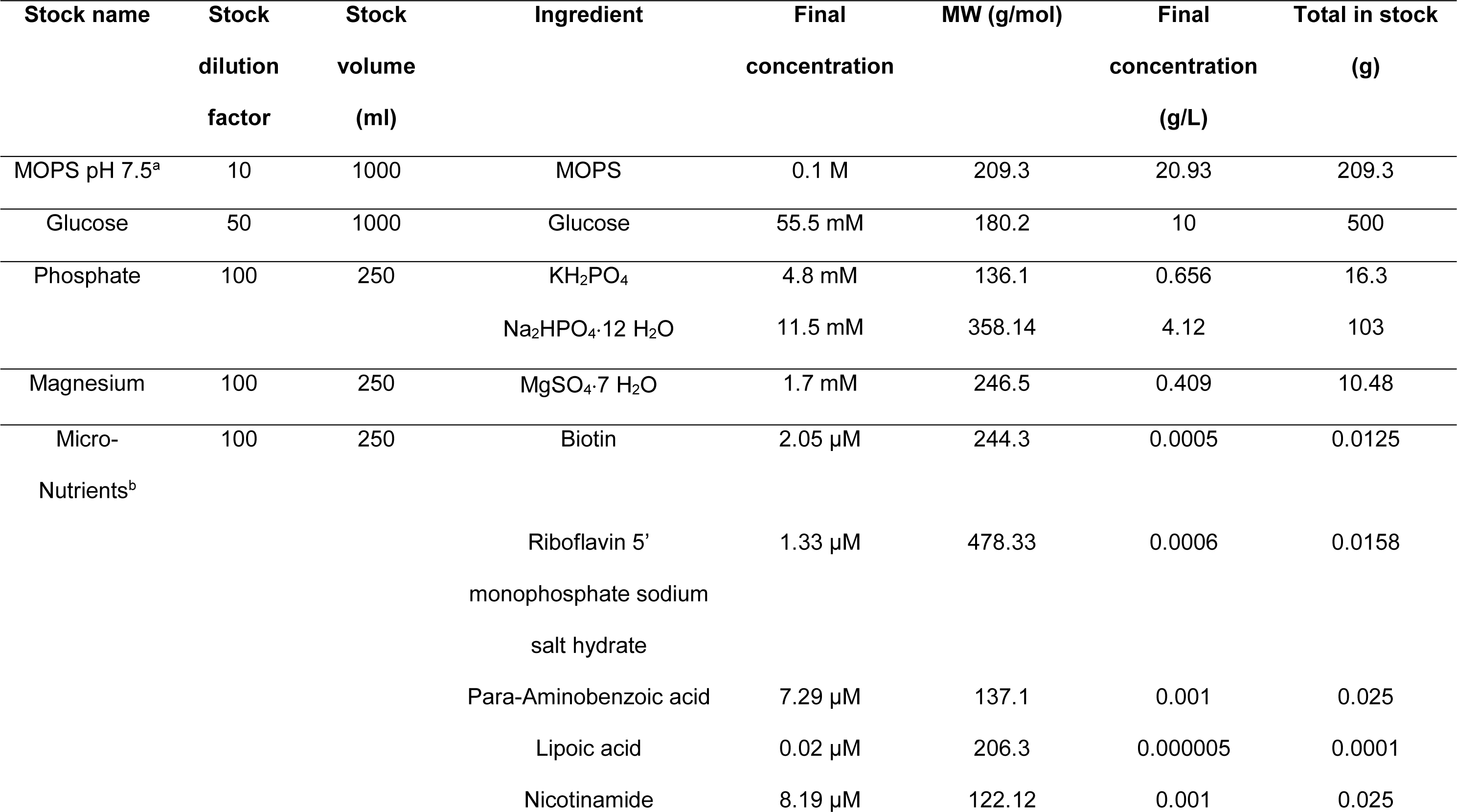
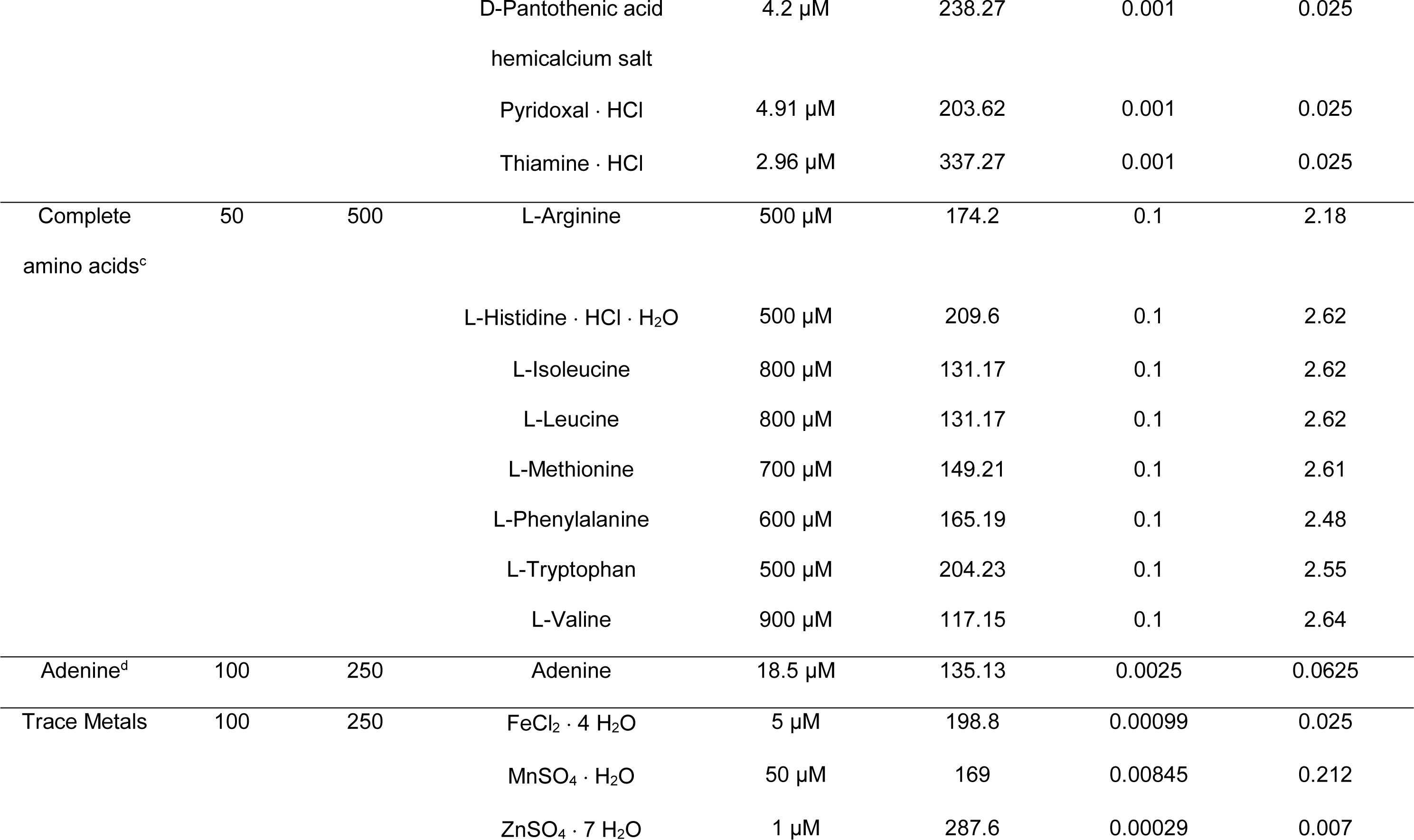
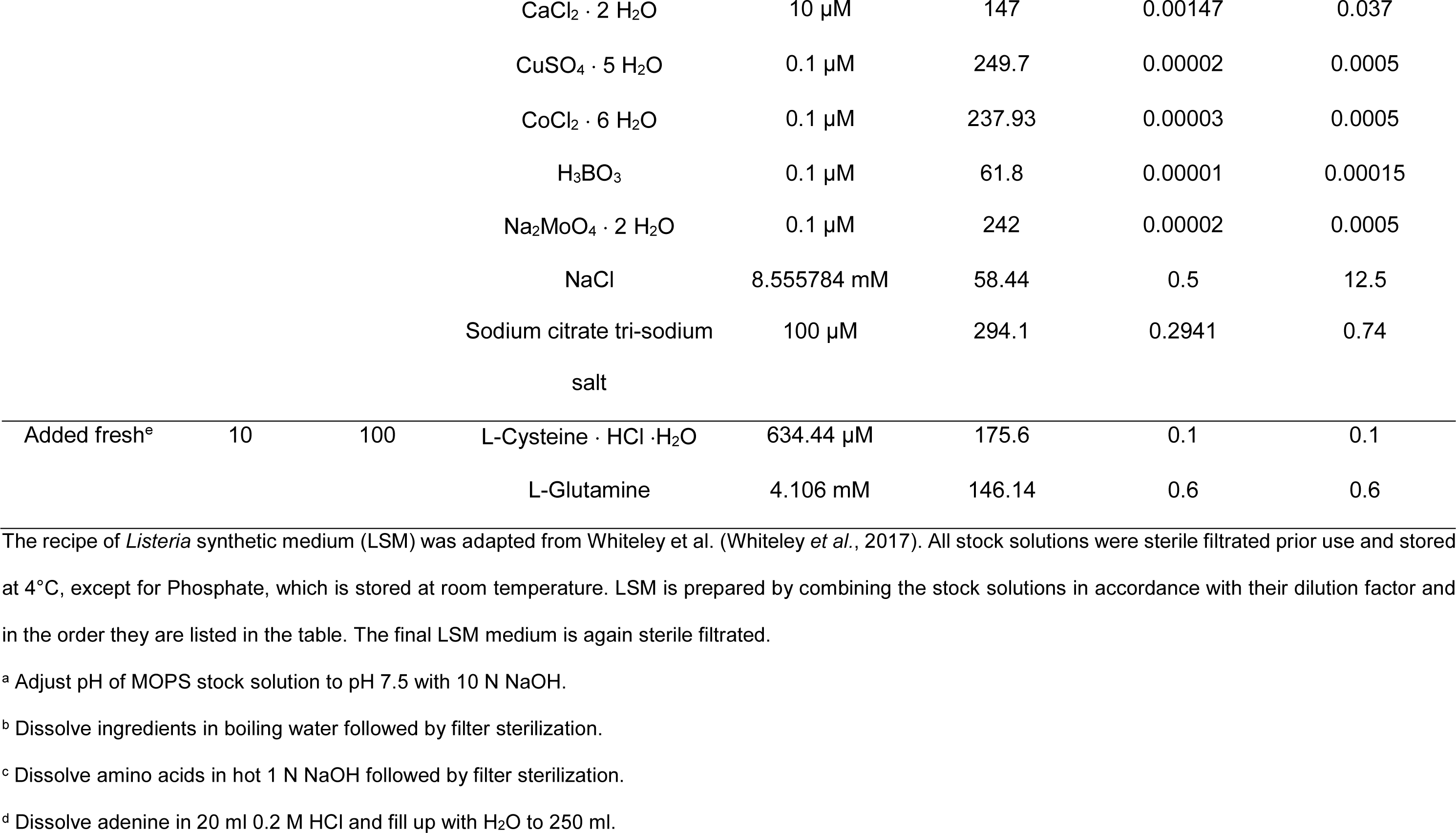
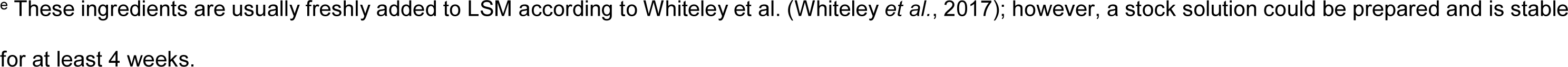
Composition of *Listeria* synthetic medium

*monocytogenes* to metabolize different carbon sources, 1% glucose, which is usually contained in LSM, was replaced by 1% of the following carbon sources: glycerol, glucosamine (Glc*N*), *N*-acetylglucosamine (Glc*N*Ac), *N*-acetylmannosamine (Man*N*Ac), glucose-1- phosphate (Glc-1-P), glucose-6-phosphate (Glc-6-P), mannose, maltose, mannitol, sucrose, rhamnose, trehalose, succinate, cellobiose, galactose and *N*-acetylneuraminic acid (Neu5Ac).

A previous study showed that the expression of *prfA* is induced in the presence of low levels of branched chain amino acids (Lobel *et al*., 2015). For growth of *L. monocytogenes* under PrfA-activating conditions, the concentration of branched chain amino acids was 10-fold reduced in freshly prepared LSM yielding 80 µM L-isoleucine, 80 µM L-leucine and 90 µM L-valine.

### Growth assays

Overnight cultures of the indicated *L. monocytogenes* strains were diluted in fresh BHI medium to an OD_600_ of 0.1 and incubated at 37°C until an OD_600_ of 0.3 was reached. Cells of 2 ml culture were collected per condition, washed twice with 1 ml of the corresponding LSM and re-suspended in 1 ml LSM. The cultures were adjusted to an OD_600_ of 0.2 and 100 µl of the cell suspension were transferred into wells of a 96-well plate containing 100 µl of the corresponding LSM. The plate was incubated at 37°C with orbital shaking in the BioTek Epoch 2 microplate reader and the OD_600_ was measured every 15 min for at least 36 h. Averages and standard deviations of at least three independent growth assays were plotted.

To test whether growth of *L. monocytogenes* was possible in LSM Glc-1-P and Glc-6- P after pre-growth of the indicated strains under PrfA-activating conditions, overnight cultures of the indicated *L. monocytogenes* strains were diluted in LSM glucose with low concentrations of BCAA to an OD_600_ of 0.1 and incubated at 37°C until an OD_600_ of 0.3 was reached. Cells were collected, washed and used to inoculate a 96-well plate containing the corresponding LSM with low concentrations of BCAA as described above.

### Strain and plasmid construction

All primers used in this study are listed in Table 3. For markerless deletion of *lmo2795*, 1-kb fragments up- and downstream of *lmo2795* were amplified by PCR using primer pairs LMS346/LMS347 and LMS348/LMS349, respectively. The resulting PCR fragments were fused by PCR using primers LMS346/LMS349. The deletion fragment was subsequently digested with *Xba*I and *Sac*I and ligated into plasmid pKSV7 that had been cut with the same enzymes. Plasmid pKSV7-Δ*lmo2795* was recovered in *E. coli* XL1-Blue yielding strain EJR88. For markerless deletion of *lmo2796*, *lmo2799*, *lmo2800* and *nanE*, 1-kb fragments up- and downstream of the corresponding gene were amplified by PCR using primer pairs LMS395/LMS396 and LMS397/LMS398 (*lmo2796*), LMS411/LMS412 and LMS413/LMS414 (*lmo2799*), JR211/LMS408 and LMS409/LMS410 (*lmo2800*), LMS401/LMS402 and LMS403/LMS404 (*nanE*). The resulting PCR fragments were fused by PCR using primers LMS395/LMS398 (*lmo2796*), LMS411/414 (*lmo2799*), JR211/LMS410 (*lmo2800*) and LMS401/LMS404 (*nanE*). The Δ*lmo2799*, Δ*lmo2800* and Δ*nanE* deletion fragments were digested with *BamH*I and *Pst*I and ligated into *BamH*I/*Pst*I cut pKSV7. Plasmids pKSV7-Δ*lmo2796,* pKSV7-Δ*lmo2799* and pKSV7-Δ*nanE* were recovered in *E. coli* XL10-Gold yielding strains EJR238, EJR122 and EJR210, respectively. pKSV7-Δ*lmo2800* was recovered in *E. coli* XL1-Blue yielding strain EJR121. For the construction of pKSV7-Δ*lmo2798* and pKSV7-Δ*lmo2797*, 1-kb fragments up- and downstream of the corresponding gene were amplified by PCR using primer pairs LMS417/LMS418 and LMS419/LMS420 (*lmo2798*) and JR142/JR143 and JR144/JR145 (*lmo2797*). The resulting PCR fragments were fused by PCR using primers LMS417/LMS420 (*lmo2798*) and JR142/JR145 (*lmo2797*), cut with *BamH*I and *EcoR*I and ligated into *BamH*I/*EcoR*I cut pKSV7. pKSV7-Δ*lmo2798* was transformed into *E. coli* XL10-Gold yielding strain EJR123. Plasmid pKSV7-Δ*lmo2797* was recovered in *E. coli* XL1-Blue yielding strain EJR164. Plasmids pKSV7-Δ*lmo2795*, pKSV7-Δ*lmo2796*, pKSV7-Δ*lmo2797*, pKSV7-Δ*lmo2798*, pKSV7-Δ*lmo2799*, pKSV7-Δ*lmo2800* and pKSV7-Δ*nanE* were transformed into *L. monocytogenes* strain EGD-e and genes *lmo2795*, *lmo2797*, *lmo2798*, *lmo2799*, *lmo2800* and *nanE* deleted by allelic exchange as previously described (Camilli *et al*., 1993) yielding strains EGD-e Δ*lmo2795* (LJR123), Δ*lmo2796* (LJR264), EGD-e Δ*lmo2797* (LJR204), EGD-e Δ*lmo2798* (LJR132), EGD-e Δ*lmo2799* (LJR135), EGD-e Δ*lmo2800* (LJR206) and EGD-e Δ*nanE* (LJR207). To construct strain EGD-e Δ*lmo2799* Δ*lmo2797* (LJR205), pKSV7-Δ*lmo2797* was transformed into *L. monocytogenes* strain LJR204 and *lmo2797* deleted by allelic exchange.

**Table 3:**
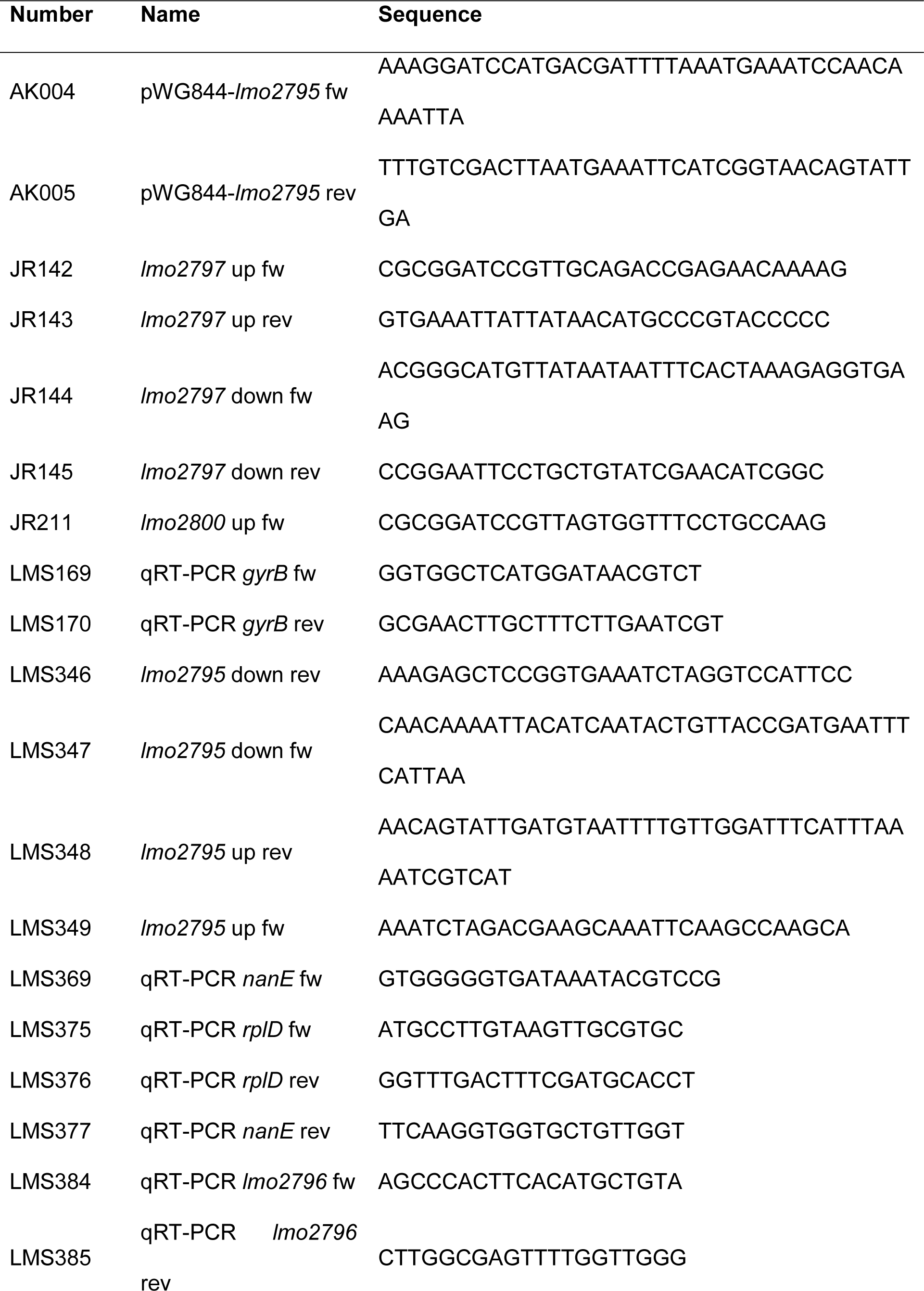
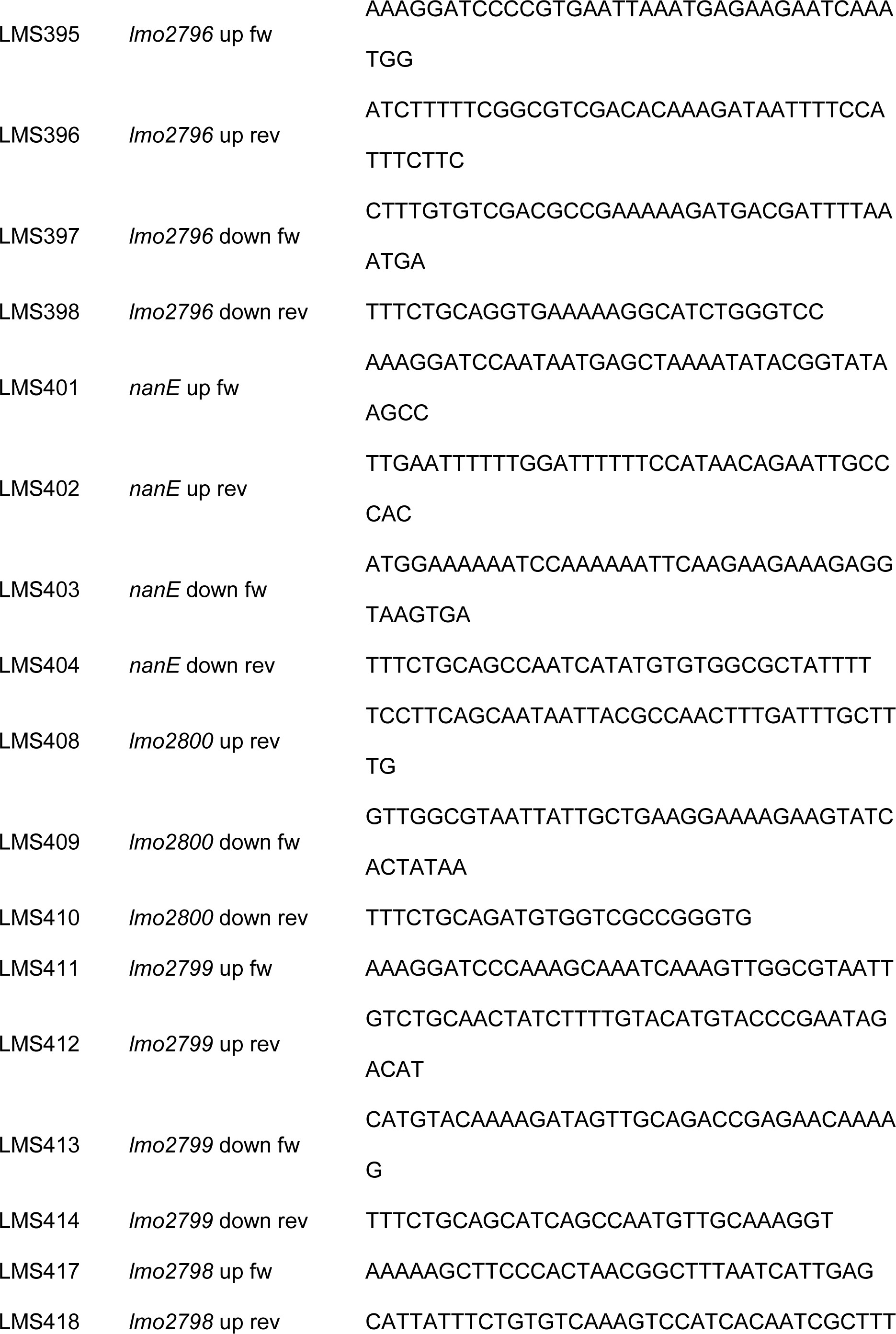
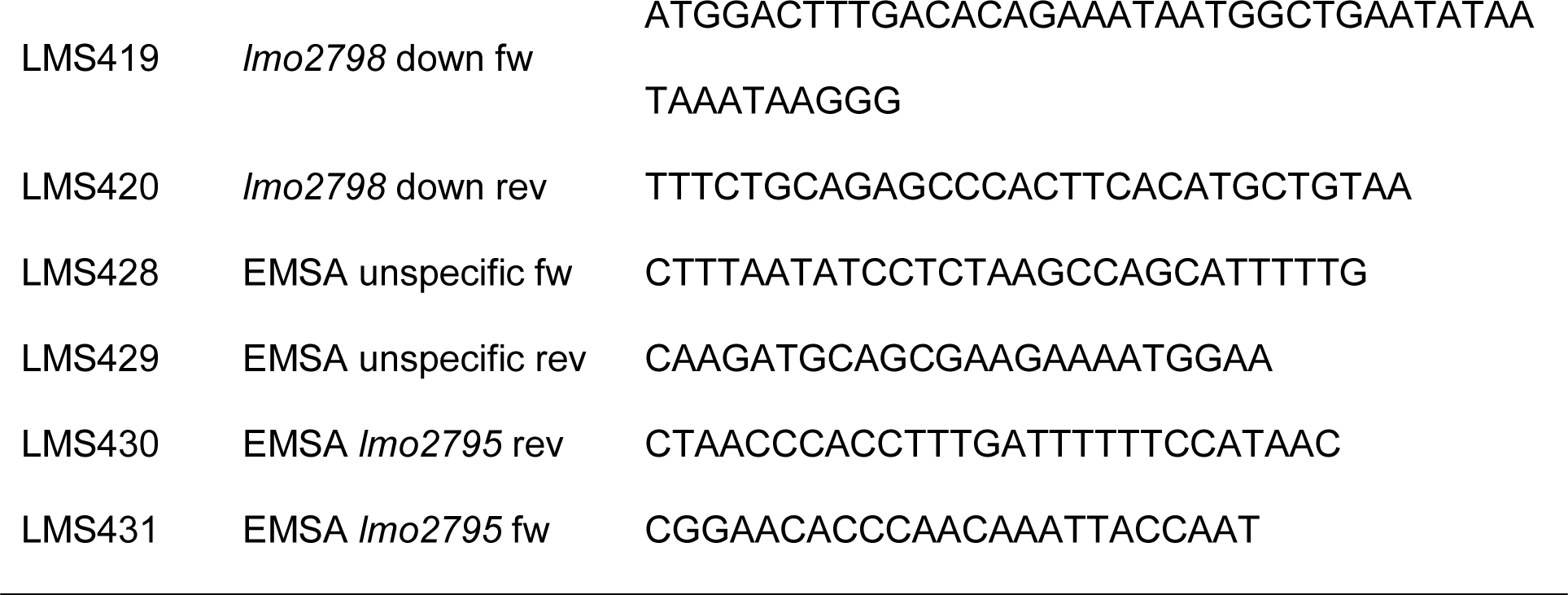
Primers used in this study

For the purification of Lmo2795 with an N-terminal His-tag, *lmo2795* was amplified using primer pair AK004/AK005. The resulting PCR product was digested with *BamH*I and *Sal*I and ligated into pWH844. Plasmid pWH844-*lmo2795* was recovered in *E. coli* DH5α yielding strain EJR131.

### RNA extraction

RNA of the indicated *L. monocytogenes* strains was isolated following a previously published method with minor modifications (Hauf *et al*., 2019). Briefly, a single colony was used to inoculate 10 ml LSM medium with 1% glucose as sole carbon source and cultures were grown overnight at 37°C. Next day, cultures were used to inoculate 30 ml of fresh LSM medium with 1% glucose to an OD_600_ of 0.1. When the cultures reached an OD_600_ of 0.5 ± 0.05, 25 ml of the cell suspensions were harvested by centrifugation for 15 min at 4,000 rpm and 4°C and the pellet was snap frozen in liquid nitrogen and stored at -80°C. To isolate the RNA, pellets were re-suspended in 1 ml killing buffer (20 mM Tris, pH 7.5, 5 mM MgCl_2_, 20 mM NaN_3_), transferred to 1.5 ml microtubes and centrifuged at 13,000 rpm for 60 sec. Cells were then re-suspended in 1 ml lysis buffer I (25 % sucrose; 20 mM Tris-HCl pH 8, 0.25 mM EDTA) and 2 µl lysozyme (100 mg ml^-1^) and incubated for 5 min on ice, followed by centrifugation at 5,000 rpm and 4°C for 5 min. Pellets were then re-suspended in 300 µl lysis buffer II (3 mM EDTA; 200 mM NaCl) and added to pre-heated (95°C) lysis Buffer III (3 mM EDTA; 200 mM NaCl; 1 % SDS). Samples were incubated for exactly 5 min at 95°C and 600 µl phenol/chloroform/isoamylalcohol (25:24:1) (PCI) was added. After shaking the samples at 700 rpm for 5 min, the two phases were separated by centrifugation at 13,000 rpm for 5 min and the upper aqueous phase was transferred to a new 1.5 ml microtube containing 600 µl PCI. The extraction was repeated with 600 µl chloroform/isoamylalcohol (25:1). Finally, the upper phase was transferred into an RNAse free tube and RNA was precipitated by addition of 0.1 x volume 3 M sodium acetate (pH 5.2) and 1.5 volumes 96 % ethanol and incubated at 20°C overnight. To precipitate the RNA, samples were centrifuged at 13,000 rpm for 15 min, washed in 70 % ethanol and dried under the fume hood. Finally, samples were re-suspended in 25 µl RNAse free H_2_O.

To avoid DNA contamination, 5 µg isolated RNA were digested with 5 µl DNase I (1 U µl^-1^, Thermo Scientific) for 40 min at 37°C. The reaction was stopped by addition of 2.5 µl 25 mM EDTA and incubation at 65°C for 10 min. To verify that the isolated RNA is free of DNA, a check PCR was performed using primers LMS169 and LMS170. Genomic DNA from *L. monocytogenes* EGD-e was used as a control.

### Quantitative real-time PCR (qRT-PCR)

qRT-PCR was carried out using the One-Step reverse transcription PCR kit, the Bio-Rad iCycler and the Bio-Rad iQ5 software (Bio-Rad, Munich, Germany). Three biological replicates were performed. Primer pairs LMS375/LMS376 and LMS169/170 were used to determine the transcript amounts of *rpID* and *gyrB*, respectively, which were used as internal controls. Transcript amounts for *lmo2796* and *nanE* were monitored using primer pairs LMS384/LMS385 and LMS369/LMS377, respectively. The average of the cycle threshold (C_T_) values of *rpID* and *gyrB* were used to normalize the C_T_- values obtained for *lmo2796* and *nanE*. For each strain, the fold changes of *lmo2796* and *nanE* expression were calculated using the ΔΔC_T_-method.

### Purification of His-Lmo2795

For the overexpression of His-Lmo2795, plasmid pWH844 was used, which contains the *lacI* gene, enabling the strictly IPTG-dependent overproduction of His-tagged proteins (Schirmer *et al*., 1997). pWH844-*lmo2795* was transformed into *E. coli* BL21 and grown in 2x LB. When cultures reached an OD_600_ of 0.6-0.8, expression was induced by the addition of IPTG at a final concentration of 1 mM and the cells were grown for 2 h at 37°C. Cells were collected by centrifugation, washed once with 1x ZAP buffer (50 mM Tris-HCl, pH 7.5, 200 mM NaCl) and the cell pellet stored at -20°C. The next day, cells were re- suspended in 1x ZAP (50 mM Tris-HCl, pH 7.5, 200 mM NaCl) and lysed by three passages (18,000 lb/in2) through an HTU DIGI-F press (G. Heinemann, Germany). The resulting crude extract was centrifuged at 46,400x g for 60 min followed by protein purification using a Ni^2+^ nitrilotriacetic acid column (IBA, Göttingen, Germany). His-Lmo2795 was eluted using imidazole and elution fractions analyzed using SDS-PAGE. Selected elution fractions were subsequently subjected to dialysis against 1x ZAP buffer at 4°C overnight. The protein concentration was determined according to the method of Bradford (Bradford, 1976) using the Bio-Rad protein assay dye reagent concentrate. Bovine serum albumin was used as standard. The protein samples were stored at -80°C until further use.

### Electrophoretic mobility shift assay (EMSA)

EMSA was performed as previously described with some modifications (Dhiman *et al*., 2014). Briefly, the upstream region of the *lmo2795*- *nanE* operon containing a putative promoter site was amplified using primer pair LMS430/LMS431. A 204 bp region within the operon, which served as an unspecific control, was amplified using primers LMS428/LMS429. Binding reactions were performed at 25°C for 20 min in 20 µl binding buffer containing 10 mM Tris-HCl, pH 7.5, 50 mM NaCl, 10% glycerol, 5 mM EDTA, 20 mM DTT and varying amounts of His-Lmo2795 (3.5 – 14 µg), as well as 250 pmol of DNA. To ease loading of the samples into the wells, 1 µl DNA loading dye was added to each sample. A 60-min pre-run was carried out at 70 V prior to sample loading. The samples were separated on an 8% polyacrylamide native gel in 0.5x TBE buffer (50 mM Tris-HCl, pH 10, 50 mM boric acid, 1 mM Na_2_EDTA) at 4°C for 2.5 h at 50 V. The gel was stained in 50 ml 0.5x TBE buffer containing 5 µl HDGreen^TM^ Plus DNA dye (INTAS, Göttingen, Germany) for 3 min, followed by a 5 min washing step in 0.5x TBE buffer, three 20 sec washing steps in H_2_O and an additional 30 min washing step in H_2_O. DNA bands were visualized using a GelDoc^TM^ XR+ (BioRad, Munich, Germany).

## Results and Discussion

### Growth of *L. monocytogenes* wildtype strains under salt and pH stress

For most physiological studies, which aimed at the characterization of salt and pH resistance of *L. monocytogenes*, bacteria were grown in complex media such as BHI or TSB. In nature as well as within the host, *L. monocytogenes* likely encounters limited access to nutrients, such as carbon and nitrogen sources or vitamins, which likely affects their ability to adapt to environmental stress conditions. We therefore assessed the ability of the two widely used *L. monocytogenes* wildtype strains EGD-e and 10403S to grow under high salt and low pH stress in the chemically defined medium LSM. Growth of EGD-e was barely affected in the presence of 1% NaCl. In the presence of 2, 3 and 4 % NaCl, growth of EGD-e was delayed as compared to LSM. At a concentration of 5% salt, EGD-e was still able to grow, but only reached an optical density of around 0.3 after 36 h. The *L. monocytogenes* wildtype strain 10403S showed a lower salt resistance as compared to EGD-e. Growth of 10403S was already impaired in LSM containing 3% salt. In the presence of 4% and 5% salt, 10403S could only reach an optical density of around 0.2 after 36 h (Fig. 1A). In contrast, *L. monocytogenes* is able to grow in the presence of up to 10% salt, when a complex medium is used (Patchett *et al*., 1992; Vasseur *et al*., 2001). To assess the pH resistance, we compared the growth of *L. monocytogenes* strains grown in standard LSM, which has a pH of 8.7, and LSM adjusted to a pH of 8, 7, 6 and 5.5. Growth of the *L. monocytogenes* wildtype strain EGD-e was similar in LSM with a pH of 7 as compared to standard LSM, while growth of 10403S was slightly reduced. Both strains had a severe growth defect in LSM with a pH of 6 and were unable to grow in LSM with a pH of 5.5 (Fig. 1B). Overall, we observed that the *L. monocytogenes* wildtype strain 10403S was less tolerant to salt and pH stress than EGD-e (Fig. 1). Interestingly, 10403S grew better in BHI medium, which was adjusted to a low pH, than EGD-e (Cheng *et al*., 2015). Resistance of *L. monocytogenes* to pH stress is mainly conferred by the activity of the glutamate decarboxylase activity (GAD) system, a process, which depends on the availability of extracellular glutamate (Cotter *et al*., 2001; Feehily *et al*., 2014). The two *L. monocytogenes* wildtype strains EGD-e and 10403S seem to use two distinct GAD systems, which confer a different degree of acid resistance. The GAD system used by 10403S was shown to give a higher acid tolerance (Feehily *et al*., 2014; Cheng *et al*., 2015). However, complex medium such as BHI likely contains higher glutamate levels than LSM, which can be used by the GAD system. Thus, the GAD system of 10403S might be less efficient when the cells are grown in LSM and are therefore less resistant to acid stress.

**Figure 1:**
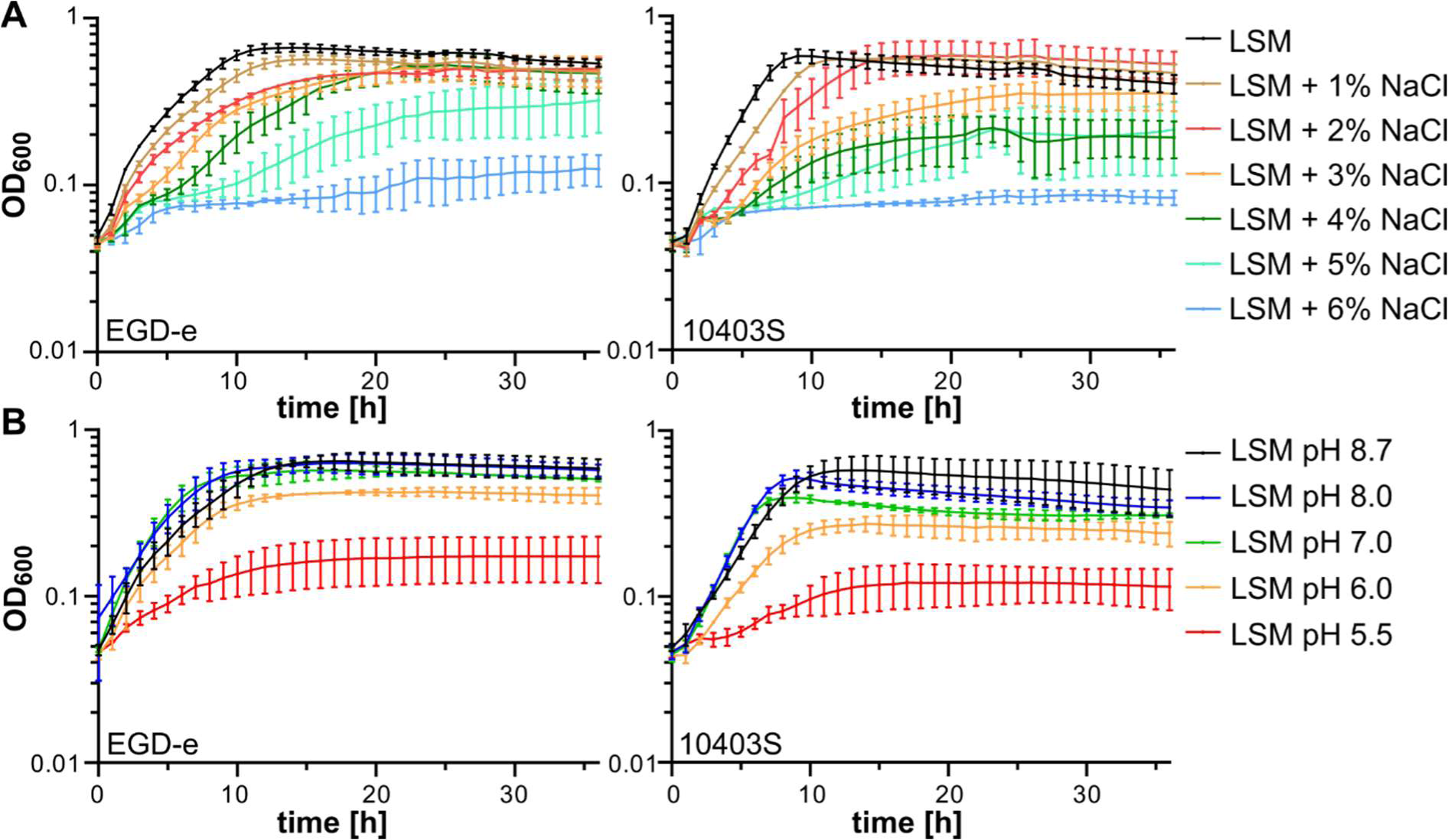
Salt and pH resistance of *L. monocytogenes* wildtype strains. (A) *L. monocytogenes* strains EGD-e and 10403S were grown in LSM containing 1% glucose and increasing concentrations of NaCl as described in the methods section. (B) *L. monocytogenes* strains EGD-e and 10403S were grown in LSM containing 1% glucose (pH 8.7). The pH of LSM was adjusted to the indicated pH-values using HCl. The average values and standard deviations of three independent experiments were plotted.

### Growth of *L. monocytogenes* in LSM with different carbon sources

Over the last decades, several chemically defined media have been developed that support growth of *L. monocytogenes* (Pine *et al*., 1989; Premaratne *et al*., 1991; Phan-Thanh and Gormon, 1997; Tsai and Hodgson, 2003; Jarvis *et al*., 2016; Whiteley *et al*., 2017). Most of these media contain glucose as the sole carbon source. Other carbon sources such as fructose and mannose were shown to support growth of *L. monocytogenes* in Hsiang-Ning Tsai medium (HTM) or modified Welshimer’s Broth (MWB) (Premaratne *et al*., 1991; Tsai and Hodgson, 2003). Glucose could also be replaced by the amino sugars Glc*N*Ac and *N*-acetylmuramic acid in MWB (Premaratne *et al*., 1991). Interestingly, lactose and rhamnose could support growth of *L. monocytogenes* in ACES-buffered chemically defined (ABCD) medium, but not in HTM and/or MWB (Table 4) (Pine *et al*., 1989; Premaratne *et al*., 1991; Tsai and Hodgson, 2003). Similarly, glycerol was shown to support growth of *L. monocytogenes* in HTM medium, while only weak growth could be observed for MWB with glycerol as sole carbon source (Table 4) (Premaratne *et al*., 1991; Tsai and Hodgson, 2003). To our knowledge, no information is available on the carbon sources that support growth of *L. monocytogenes* in LSM. We therefore performed growth experiments with *L. monocytogenes* wildtype strains EGD-e and 10403S in LSM containing 1% of selected carbon sources. This analysis revealed that growth of both strains was similar in LSM containing Glc*N*Ac, Glc*N*, mannose, cellobiose or trehalose as sole carbon source compared to glucose (Fig. 2, Table 4). Both strains were also able to use glycerol as carbon sources, while only weak growth was obtained for maltose, rhamnose and succinate (Fig. 2, Table 4). No growth was observed for both strains, when glucose was replaced with galactose, sucrose, glucose-1-phosphate (Glc-1-P), glucose-6-phosphate (Glc- 6-P) or mannitol (Fig. 3A, Table 4). The hexose phosphate transporter Hpt, which is required for the import of Glc-1-P and Glc-6-P, is expressed in a PrfA-dependent manner. The expression and activity of PrfA is controlled on transcriptional, translational and post- translational level (reviewed in (Xayarath and Freitag, 2012; Gaballa *et al*., 2019)). One factor involved in the control of *prfA* expression is the global regulator CodY, which senses the presence of branched chain amino acids (BCAAs). In the presence of high levels of BCAAs, CodY acts as a *prfA* repressor, while expression of *prfA* is induced when BCAA levels drop, which occurs upon invasion of host cells (Lobel *et al*., 2015). We therefore tested whether reduction of the BCAA concentration in LSM could lead to growth of *L. monocytogenes* wildtype strains EGD-e and 10403S on Glc-1-P and Glc-6-P as sole carbon source. While both strains grew fine in LSM with low concentrations of BCAA and glucose as sole carbon source, no growth was observed in LSM with Glc-1-P or Glc-6-P (Fig. 3A). For growth analysis, *L. monocytogenes* strains were pre-grown in BHI complex medium, in which PrfA is inactive (Renzoni *et al*., 1997). We thus wondered whether the strains have simply not sufficient time to activate PrfA and by this also do not induce the expression of *hpt*, encoding the Glc-1-P and Glc-6-P importer, when they are transferred into LSM containing the hexose phosphate sugars. To test this, we pre-grew *L. monocytogenes* strains EGD-e and 10403S in LSM glucose with low levels of BCAA and subsequently transferred them either into the same medium as a control, or into LSM with low levels of BCAA and containing Glc-1-P or Glc-6-P as sole carbon source. A strain lacking the virulence regulator PrfA was used as a control. Indeed, EGD-e and 10403S were now able to metabolize Glc-1-P and Glc-6-P, while the *prfA* mutant was unable to grow (Fig. 3B). In contrast, absence of PrfA results in a growth advantage in LSM with glucose as sole carbon source, suggesting that the expression and/or activity of PrfA is a burden for *L. monocytogenes*, which has previously been shown (Vasanthakrishnan *et al*., 2015; Friedman *et al*., 2017). Therefore, characterization of growth and the general physiology of *L. monocytogenes* in the presence of Glc-1-P and Glc-6-P as sole carbon source, requires a pre-growth under PrfA-activating conditions, for instance in a chemically defined medium or in BHI medium containing activated charcoal, Chelex or Amberlite XAD (Chico-Calero *et al*., 2002; Portman *et al*., 2017; Gaballa *et al*., 2021).

**Figure 2:**
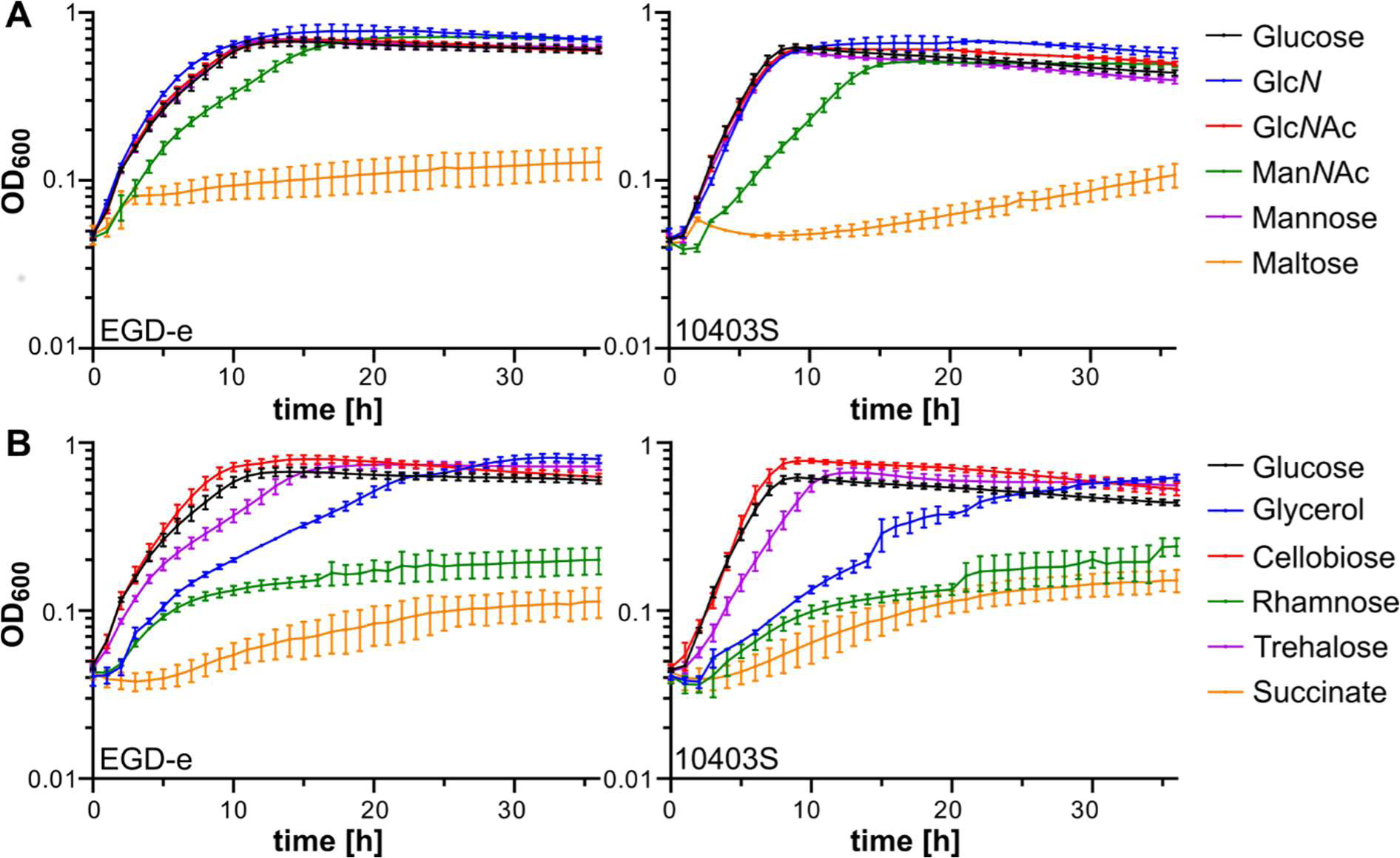
Growth in LSM media with different carbon sources. (A-B) *L. monocytogenes* strains EGD-e and 10403S were grown in *Listeria* synthetic medium (LSM) containing 1% of the indicated carbon sources as described in the methods section. The average values and standard deviations of three independent experiments were plotted. Glc*N*Ac = *N*- acetylglucosamine, Man*N*Ac = *N*-acetylmannosamine, Glc*N* = glucosamine.

**Figure 3:**
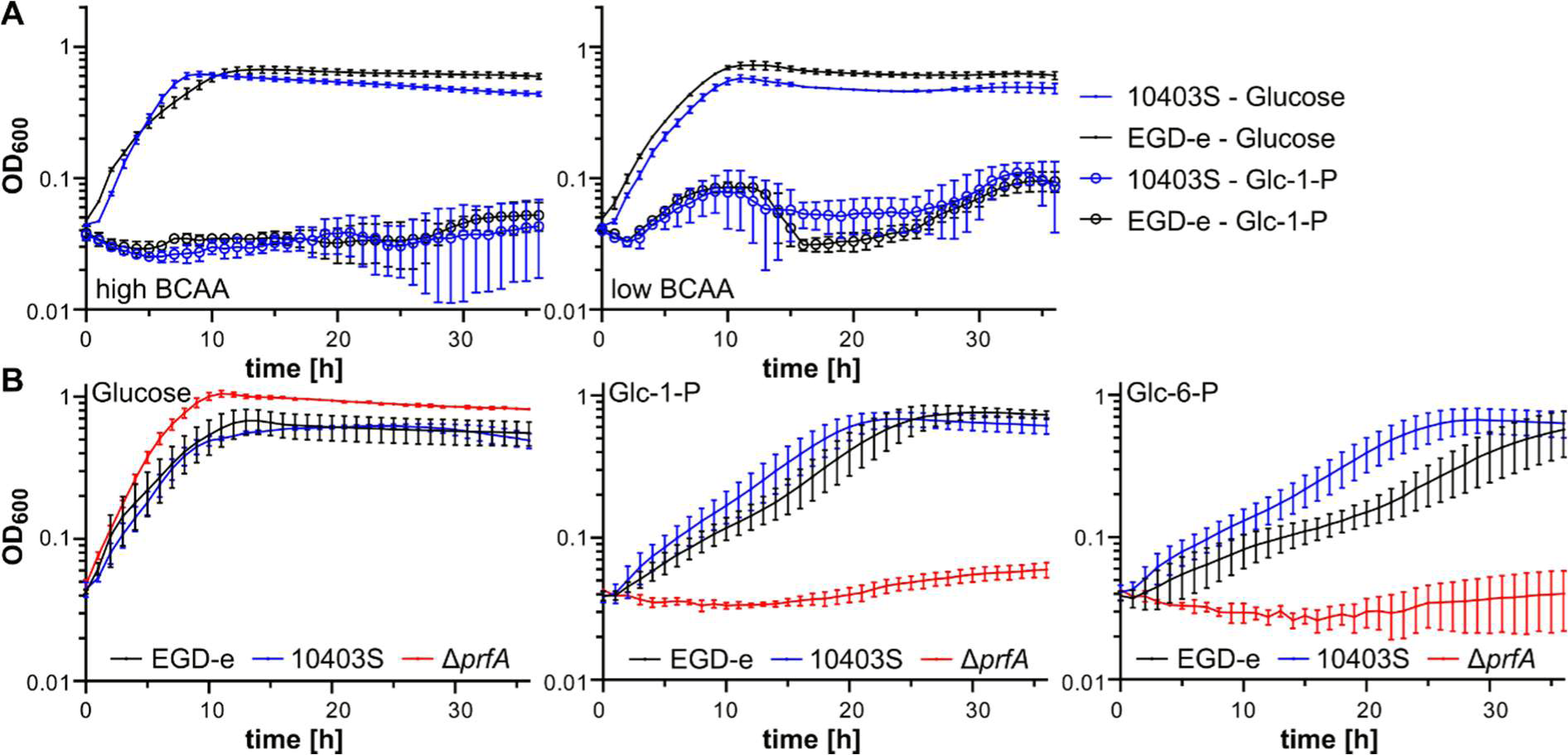
Pre-growth of *L. monocytogenes* in LSM glucose enables growth on glucose- 1-phosphate and glucose-6-phosphate. (A) *L. monocytogenes* strains EGD-e and 10403S were grown in *Listeria* synthetic medium (LSM) containing 1% of the indicated carbon sources as described in the methods section. LSM was either prepared with the standard complete amino acid mix (high BCAA) or the complete amino acid mix with 10-fold less branched chain amino acids (low BCAA). The average values and standard deviations of three independent experiments were plotted. (B) *L. monocytogenes* strains EGD-e, 10403S and Δ*prfA* were pre- grown in *Listeria* synthetic medium (LSM) containing 1% glucose and subsequently transferred to LSM containing the indicated carbon sources as described in the methods section. The average values and standard deviations of four independent experiments were plotted. Glc-1- P = glucose-1-phosphate, Glc-6-P = glucose-6-phosphate.

**Table 4:** Carbon sources supporting growth of *L. monocytogenes* in chemically defined media.

|  | (Pine *et al.*, 1989) |  | (Tsai and Hodgson, 2003) |  | (Premaratne *et al.*, 1991) |  | This study |  |
| --- | --- | --- | --- | --- | --- | --- | --- | --- |
|  | ABCD medium<sup>3</sup> |  | HTM<sup>3</sup> |  | MWB<sup>3</sup> |  | LSM<sup>3</sup> |  |
| Carbon source<sup>1</sup> | Conc.<sup>2</sup> | Growth | Conc.<sup>2</sup> | Growth | Conc.<sup>2</sup> | Growth | Conc.<sup>2</sup> | Growth |
| Glucose | 0.25% | yes | 1% | yes | 1% | yes | 1% | yes |
| Fructose |  |  | 1% | yes | 1% | yes |  |  |
| Galactose | 0.25% | no | 1% | no | 1% | no | 1% | no |
| Mannose |  |  | 1% | yes | 1% | yes | 1% | yes |
| Mannitol |  |  | 1% | no | 1% | no | 1% | no |
| Arabinose |  |  | 1% | no | 1% | no |  |  |
| Ribose |  |  | 1% | no | 1% | no |  |  |
| Xylose | 0.25% | no | 1% | no | 1% | no |  |  |
| Lactose | 0.25% | yes | 1% | no | 1% | no |  |  |
| Rhamnose | 0.25% | yes | 1% | no |  |  | 1% | weak |
| Maltose |  |  | 1% | no | 1% | weak | 1% | weak |
| Sucrose |  |  | 1% | no | 1% | no | 1% | no |
| Cellobiose |  |  |  |  | 1% | yes | 1% | yes |
| Trehalose |  |  |  |  | 1% | yes | 1% | yes |
| Glc-1-P |  |  |  |  | 1% | no | 1% | yes<sup>4</sup> |
| Glc-6-P |  |  |  |  | 1% | no | 1% | yes<sup>4</sup> |
| GlcN |  |  |  |  | 1% | yes | 1% | yes |
| GlcNAc |  |  |  |  | 1% | yes | 1% | yes |
| MurNAc |  |  |  |  | 1% | yes |  |  |
| ManNAc |  |  |  |  |  |  | 1% | yes |
| Neu5Ac |  |  |  |  |  |  | 1% | no |
| Glycerol |  |  | 1% | yes | 1% | weak | 1% | yes |
| Succinate |  |  |  |  | 1% | no | 1% | weak |

<sup>1</sup> The following carbon sources do not support growth: Melibiose, Raffinose, Sorbose, Sorbitol, Galactose-1-phosphate, Galactose-6-phosphate, Lactate, Citrate, Isocitrate, $\alpha$ -Ketoglutarate, Fumarate, Malate, Pyruvate, Acetate, Chitin (Premaratne *et al.*, 1991). Abbreviations: Glc-1-P = Glucose-1-phosphate, Glc-6-P = Glucose-6-phosphate, GlcN = glucosamine, GlcNAc = *N*-acetylglucosamine, MurNAc = *N*-acetylmuramic acid, ManNAc = *N*-acetylmannosamine, Neu5Ac = *N*-acetylneuraminic acid.
<sup>2</sup> Conc. = concentration of carbon source
<sup>3</sup> ABCD medium = *N*-(2-acetamido)-2-aminoethanesulfonic acid (ACES)-buffered, chemically defined medium; HTM = Hsiang-Ning Tsai medium, MWB = Modified Welshimer's broth, LSM = *Listeria* synthetic medium
<sup>4</sup> Pre-growth in LSM glucose with low concentrations of BCAA, a PrfA-activating condition, is required.

The amino sugar Man*N*Ac, which has not been tested for any of the other chemically defined media, was able to support growth of *L. monocytogenes* strains EGD-e and 10403S (Fig. 2A, Table 4). Man*N*Ac is the precursor of the sialic acid *N*-acetylneuraminic acid (Neu5Ac), which can serve as a carbon source for *Staphylococcus aureus* inside the host (Angata and Varki, 2002; Vimr *et al*., 2004; Olson *et al*., 2013). However, Neu5Ac could not support growth of *L. monocytogenes*, suggesting that this organism does not contain the necessary catabolic pathway (Table 4).

### The *lmo2795-nanE* operon is required for Man*N*Ac catabolism

In *E. coli*, Man*N*Ac is imported into the cell by the PTS transporter ManXYZ yielding Man*N*Ac- 6-phosphate. Man*N*Ac-6-phosphate is subsequently converted into Glc*N*Ac-6-phosphate by the Man*N*Ac-6-phosphate epimerase NanE. The genome of *L. monocytogenes* strain EGD-e contains one homolog of NanE encoded by *lmo2801* (37% identity, 58% similarity). Based on the presence of the NanE homolog, it was already previously proposed that the *lmo2795*-*nanE* operon might be involved in the transport of Man*N*Ac (Deutscher *et al*., 2014). In addition to NanE, the *lmo2795-nanE* operon encodes an RpiR-type regulator, a member of the repressor, ORF, kinase (ROK) family, the EIIA and EIIBC components of a PTS system, a putative HAD hydrolase and a putative oxidoreductase (Fig. 4A). To test whether these proteins are required for the catabolism of Man*N*Ac, single deletion mutants and a mutant lacking both PTS components, Lmo2797 and Lmo2799, were constructed in the EGD-e wildtype background and their growth was analyzed. All deletion mutants were able to grow in LSM containing glucose, however, growth was enhanced for *L. monocytogenes* strains Δ*lmo2799* and Δ*lmo2797* Δ*lmo2799*, lacking one or two of the PTS components, and the *nanE* deletion mutant (Fig. 4B-C). In contrast, growth on Man*N*Ac as sole carbon source was reduced for the *lmo2795* and *lmo2798* deletion strains lacking the RpiR transcriptional regulator and the putative HAD hydrolase, respectively, and was nearly abolished for the strain lacking NanE (Fig. 4D-E). Surprisingly, deletion strains Δ*lmo2799* and Δ*lmo2797* Δ*lmo2799* showed a better growth in LSM with Man*N*Ac, similar to what was observed for the glucose containing LSM (Fig. 4C-E), indicating that this PTS system is either not involved in the import of Man*N*Ac or that Man*N*Ac can also be imported by other PTS systems of *L. monocytogenes*. Absence of individual PTS systems can also lead to the overexpression of another PTS system (Stoll and Goebel, 2010), which could explain why strains lacking the EIIB/C component Lmo2799 grew better than the wildtype in LSM with glucose or Man*N*Ac.

**Figure 4:**
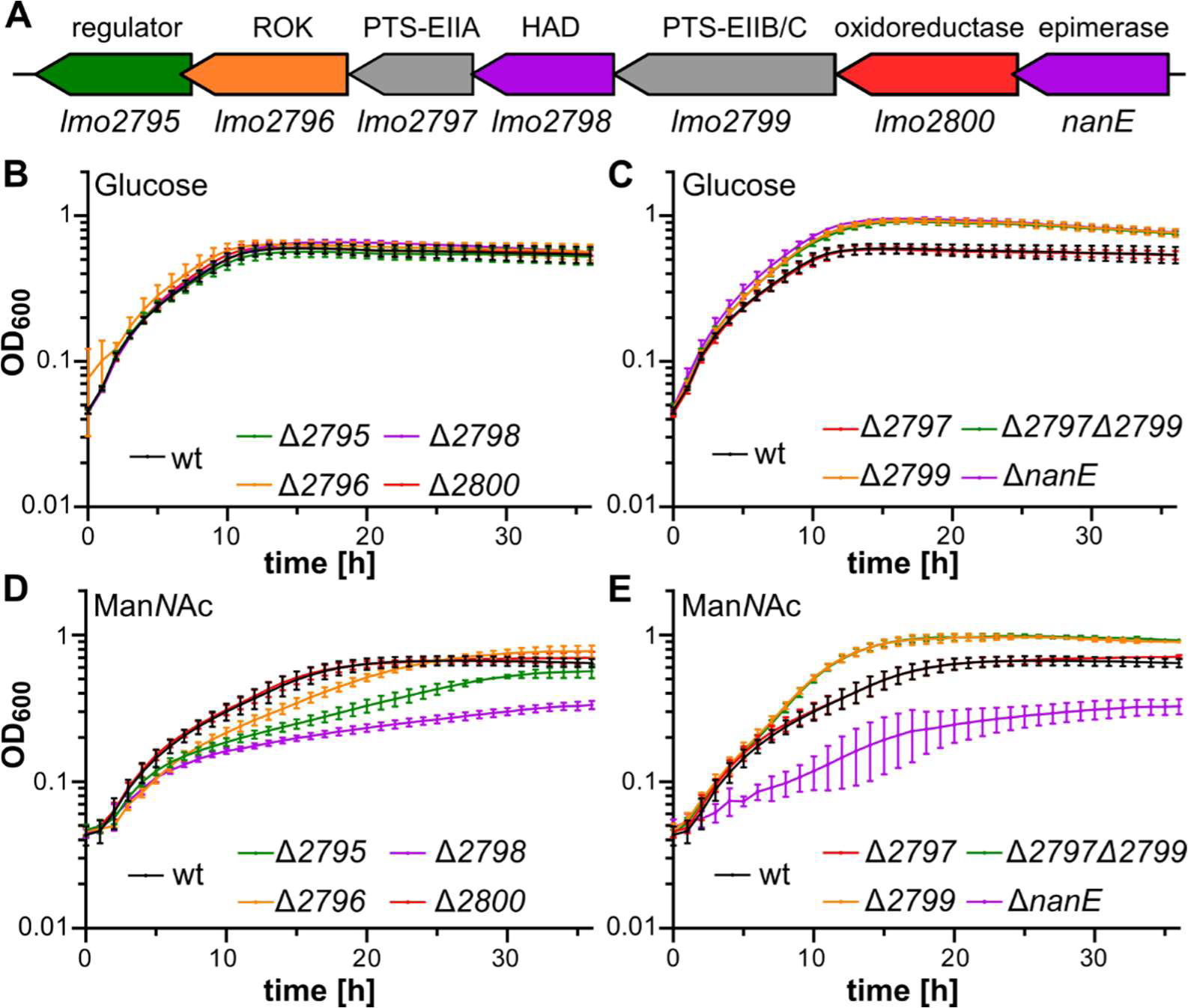
The *lmo2795-nanE* operon is required for Man*N*Ac utilization. (A) Gene arrangement of the *lmo2795-nanE* operon of *L. monocytogenes*. Predicted protein functions are indicated above the genes. ROK – member of the repressor, ORF, kinase family; PTS-EIIA and PTS-EIIB/C – components of a phosphotransferase system; HAD – member of the haloacid dehydrogenase-like hydrolase superfamily. (B-G) Growth curves. The indicated *L. monocytogenes* strains were grown in LSM containing (B, C) 1% glucose, (D, E) 1% Man*N*Ac as sole carbon source. The average values and standard deviations of three independent experiments were plotted.

### Expression of the *lmo2795-nanE* operon is controlled by Lmo2795

RpiR-type transcriptional regulators are often involved in the control of sugar metabolic pathways in Gram-positive and Gram-negative bacteria (Sørensen and Hove-Jensen, 1996; Yamamoto *et al*., 2001; Jaeger and Mayer, 2008; Li *et al*., 2017; Aleksandrzak-Piekarczyk *et al*., 2019). We thus wondered whether Lmo2795 regulates the expression of the *lmo2795*- *nanE* operon. First, we conducted EMSA experiments to assess whether Lmo2795 binds to the promoter region of the operon. For this purpose, increasing concentrations of purified His- Lmo2795 (Fig. 5A) were incubated with the DNA fragment containing the putative promoter of the *lmo2795*-*nanE* operon. The migration of the DNA fragment was retarded in the presence of His-Lmo2795 as compared to the DNA sample that did not contain the protein, suggesting that a protein-DNA complex was formed. In addition, a so-called supershift could also be observed, whose intensity increases with increasing protein concentrations. In contrast, no retardation of the unspecific DNA control was observed in the presence of His-Lmo2795 (Fig. 5B). To test whether Lmo2795 activates or represses the expression of the *lmo2795*-*nanE*, we quantified the expression of *lmo2796* and *nanE* via quantitative real-time PCR. To allow for a better comparison of the gene expression, we decided to isolate RNA of the *L. monocytogenes* wildtype strain EGD-e and the *lmo2795* mutant that were grown in LSM with glucose as sole carbon source due to the observed growth deficit of the *lmo2795* mutant in Man*N*Ac-containing LSM. This analysis revealed that the expression of both genes, *lmo2796* and *nanE*, was reduced in the *lmo2795* mutant as compared to the wildtype strain (Fig. 5C), suggesting that Lmo2795 acts as a transcriptional activator under the tested condition. Based on the observation that a strain lacking NanE had a growth deficit in LSM with Man*N*Ac as sole carbon source, we assume that Lmo2795 also activates the expression of the *lmo2795*-*nanE* under this condition as we would expect at least wildtype-like growth of the *nanE* mutant, if Lmo2795 acts as a repressor. Further analysis is required to support this hypothesis.

**Figure 5:**
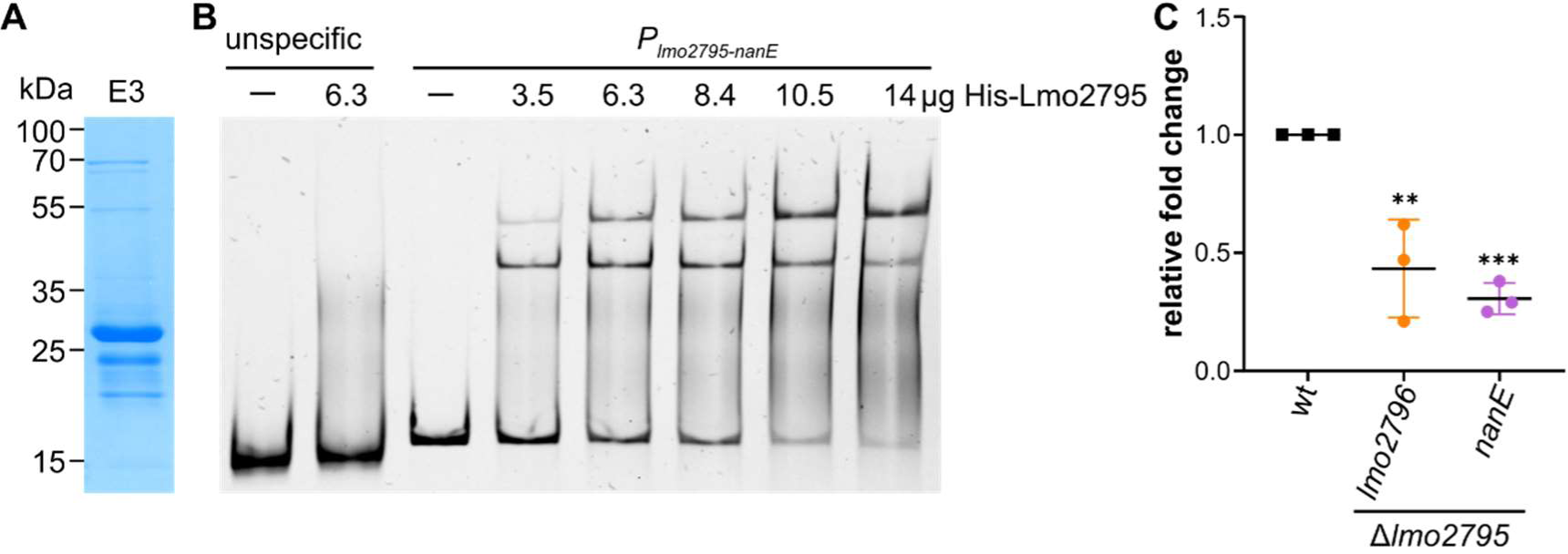
Lmo2795 regulates the expression of the *lmo2795-nanE* operon. (A) SDS-PAGE of elution fraction 3 containing purified His-Lmo2795. The expected molecular weight of His- Lmo2795 is 31.5 kDa. (B) EMSA assay. Increasing concentrations of His-Lmo2795 were incubated with the DNA fragment containing the predicted promoter region of the *lmo2795- nanE* operon. The DNA-protein complexes were separated on an 8% polyacrylamide gel, which was subsequently stained using HDGreen^TM^. A DNA fragment containing a short region from within the operon was used as the unspecific control. Reactions without protein were used as negative controls (-). (C) Analysis of *lmo2796* and *nanE* expression by qRT-PCR. RNA was isolated from *L. monocytogenes* strains EGD-e (wt) and Δ*lmo2795* grown in LSM containing 1% glucose as sole carbon source as described in the methods section. Expression of *lmo2796* and *nanE* was normalized to the expression of *gyrB* and *rplD* and fold changes were calculated using the ΔΔC_T_ method. Averages and standard deviations of three independent RNA extractions were plotted. For statistical analysis, a one-way ANOVA coupled with a Dunnett‘s multiple comparison test was performed (** *p* ≤ 0.01; *** *p* ≤ 0.001).

## Conclusion

The human pathogen *L. monocytogenes* is able to withstand and adapt to a variety of environmental stress conditions. Most studies on physiology and stress tolerance of this organism were performed in complex media without any nutrient limitation; however, this condition is rarely found in their natural habitat or within the host during infection. Our study focused on the characterization of the growth of two widely used *L. monocytogenes* wildtype strains in the newly developed *Listeria* synthetic medium. We were able to show that both strains can utilize a variety of carbon sources, when supplied as sole carbon source. To our knowledge, this is the first time that the catabolism of Man*N*Ac has been investigated for *L. monocytogenes*. We were able to show that the growth of this pathogen depends on the presence of NanE, which is encoded in the *lmo2795-nanE* operon. The expression of the *lmo2795-nanE* operon is regulated by the RpiR-type regulator Lmo2795, which acts as a transcriptional activator under the tested conditions.

Our detailed characterization of the growth of *L. monocytogenes* in LSM in the presence of diverse stress conditions or carbon sources can serve as a starting point for future studies focusing on the adaptation of this important human pathogen on changing environmental conditions.

## Acknowledgements

We thank Julia Busse for technical assistance, Fabienne Dreier and Tayfun Acar for their help with constructing strains and plasmids and Pascale Cossart for sharing the *prfA* mutant. We are grateful to Prof. Jörg Stülke for providing JR and LMS with laboratory space, equipment and consumables and to the Göttingen Center for Molecular Biosciences (GZMB) for financial support.

## Author Contributions

LMS: Investigation, formal analysis, conceptualization, methodology, supervision, validation, writing – review & editing. AK: Investigation, formal analysis, writing – review & editing. JR: Investigation, conceptualization, supervision, funding acquisition, writing – original draft.

